# Stable and dynamic gene expression patterns over diurnal and developmental timescales in *Arabidopsis thaliana*

**DOI:** 10.1101/2024.09.18.613638

**Authors:** Ethan J Redmond, James Ronald, Seth J Davis, Daphne Ezer

## Abstract

- Developmental processes are known to be circadian regulated in plants. For instance, the circadian clock regulates genes involved in the photoperiodic flowering pathway and the initiation of leaf senescence. Furthermore, signals which entrain the circadian clock, such as energy availability, are known to vary in strength over plant development. However, diurnal oscillations of the Arabidopsis transcriptome have typically been measured in seedlings.
- We collected RNA-seq data from Arabidopsis leaves over developmental and diurnal timescales, concurrently: every 4 hours per day, on 3 separate days after a synchronised vegetative-to-reproductive transition. Gene expression varied more over the developmental timescale than on the diurnal timescale, including genes related to a key energy sensor: the Sucrose non-fermenting-1-related protein kinase (SnRK1) complex.
- Moreover, regulatory targets of core clock genes displayed changes in rhythmicity and amplitude of expression over development. Cell-type-specific expression showed diurnal patterns that varied in amplitude, but not phase, over development. Some previously identified qRT-PCR housekeeping genes display undesirable levels of variation over both timescales. We identify which common qRT-PCR housekeeping genes are most stable across developmental and diurnal timescales.
- In summary, we establish the patterns of circadian transcriptional regulation over plant development, demonstrating how diurnal patterns of expression change over developmental timescales.

## Introduction

Due to their sessile nature within a cyclical environment, plants have evolved an internal timekeeper called the circadian clock. It consists of a core interlocking loop of transcriptional regulators, whose components are mostly conserved across embryophytes (Petersen *et al*., 2022; Wang *et al*., 2024). Circadian clocks confer fitness to plants by allowing them to coordinate responses to photoperiod, light quality, temperature, and other environmental cues (Dodd *et al*., 2005; Atamian *et al*., 2016; Rubin *et al*., 2017; Xu *et al*., 2022). This coordination takes place through widespread transcriptional control of the plant transcriptome, among other methods of regulation (Nagel *et al*., 2015; Hayama *et al*., 2017; Ezer *et al*., 2017; Romanowski *et al*., 2020; Xiong *et al*., 2022). Estimates of the proportion of the *Arabidopsis* and wheat transcriptomes that are controlled by the clock range from 30% to 50% (Covington *et al*., 2008; Romanowski *et al*., 2020; Rees *et al*., 2022).

Large-scale circadian RNA-seq experiments are typically performed in seedlings or juvenile plants, due to the ease of performing timeseries experiments under different entrainment and free-running conditions. Yet, many of the clock genes have fundamental roles in the timing of developmental processes (Inoue *et al*., 2018; Wang *et al*., 2024). A key example of this is the evening complex (EC), a tripartite protein complex containing EARLY FLOWERING 3 and 4 (ELF3 and ELF4) and LUX ARRHYTHMO (LUX) (Nusinow *et al*., 2011; Herrero *et al*., 2012). The EC, in combination with GIGANTEA (GI), acts upstream of the photoperiodic flowering pathway in *Arabidopsis* and rice (Fowler *et al*., 1999; Park *et al*., 1999; Sawa & Kay, 2011; Andrade *et al*., 2022). Leaf senescence occurs concurrently with the vegetative-to-reproductive transition (Redmond et al., 2024) and is also regulated by the circadian clock. Mutations in clock genes, including *ELF3, PSEUDO-RESPONSE REGULATOR9* (*PRR9*) and *CIRCADIAN-CLOCK ASSOCIATED1* (*CCA1*) have all been shown to affect the onset of leaf senescence (Sakuraba *et al*., 2014; Song *et al*., 2018; Kim *et al*., 2018). This emphasises the need to study circadian rhythms in adult plants, when the relevant developmental transitions are occurring.

Moreover, developmentally associated processes exhibit diurnal patterns. These patterns begin at the earliest stages of plant development and extend into all developmental stages. Germination responds to diurnally fluctuating temperatures in many plant species (Thompson *et al*., 1977). Hypocotyl development is mediated through auxin- and temperature-related processes, via the clock-controlled *PHYTOCHROME-INTERACTING FACTORS 4 and 5 (PIF4* and *PIF5)* (Nozue *et al*., 2007; Seaton *et al*., 2015). Many of the central integrators that control flowering time are diurnally expressed, such as *FLOWERING LOCUS T* (*FT*), *SUPRESSOR OF OVEREXPRESSION OF CO1* (*SOC1*) and *LEAFY* (*LFY*) (Wendell *et al*., 2017). Additionally, many of the transcription factors that regulate the synthesis of key plant hormones involved in development, like auxin and ABA, are diurnally expressed (Balcerowicz *et al*., 2021).

Tissue-specificity also plays a role in the link between the clock and development. For instance, the vascular clock plays a dominant role over the epidermal clock in leaves. Moreover, these tissue-specific clocks influence two distinct developmental processes, flowering and hypocotyl development (Endo *et al*., 2014). Vong *et al*., 2024 suggested that each cell type’s transcriptional activity varied across diurnal and developmental timescales, but it is unclear how the daily oscillations in cell-type activity vary over developmental timescales.

There remains a large gap in our knowledge of how diurnal genes vary over development and how developmental genes vary across the day. Here, we address this gap by measuring diurnal gene expression over the one of the most crucial developmental transitions an annual plant experiences: the vegetative-to-reproductive transition. We measure gene expression over two timescales: every 24 hours (the diurnal timescale) and over approximately 2 weeks (the developmental timescale). As individual plants experience developmental asynchrony in their floral transition (Klingenberg, 2019; Redmond *et al*., 2024), we use a photoperiod shift from short days (SD; 8h light per day) to long days (LD; 16h light per day) to induce synchronised vegetative-to-reproductive transitions. One day of LD conditions is sufficient to induce maximum gene expression response of *FLOWERING LOCUS T* (*FT*) (Corbesier *et al*., 1996; Krzymuski *et al*., 2015). We therefore measure diurnal gene expression in ageing plants after the inductive SD to LD signal.

Here, we determine the extent of transcriptional changes over both developmental and diurnal timescales. First, we find that gene expression changes most dramatically over the developmental timescale and that core clock genes have broadly stable phase and amplitude of expression per day. Second, we observe that the expression dynamics of targets of a key *Arabidopsis* energy sensor vary across both scales. Third, we show that transcriptional targets of core clock genes exhibit differing changes of amplitude over development. Fourth, we identify tissue-specific changes in rhythmic processes in ageing leaves. Finally, one of the most important applications of our work is in identifying sets of genes that are stable across both timescales, as these could serve as important controls for qRT-PCR. We suggest a filtered set of housekeepers that we would encourage the community to use for studies which aim to study gene expression of clock-controlled processes in ageing plants.

## Materials and Methods

### Plant growth conditions

Seeds from the Arabidopsis ecotype Wassilewskija (Ws-2) (Anwer *et al*., 2020) were surface-sterilised. They were then plated onto 1x Murashige and Skoog basal salts (Duchefa Biochemie) supplemented with 1% w/v sucrose (ThermoFisher), 0.5% w/v MES (Melford Bioscience) and 1.5% w/v phytoagar (Duchefa Biochemie). After four days of stratification at 4°C, these plates were then transferred to short day (SD) conditions (8h light / 16h dark; 70 µmol m^−2^ s^−1^ with vertical fluorescent lighting; Osram L 36W/830 Warm White) at 21°C and left for 10 days. Then, 2 seedlings per pot were transferred to soil, then were thinned out to 1 seedling per pot after 3 days. 3rd and 4th true rosette leaves were tracked for each individual plant, by placing pipette tips in soil next to these leaves and adjusting them as the leaves moved. 12 days after transferring to soil, the conditions were changed to long day (LD; 16h light / 8h dark) at 21°C.

Samples were collected at ZT (zeitgeiber time; hours after dawn) 0, 1, 4, 8, 12, 16, 20 on three days. We chose to include the ZT1 timepoint due to our previous observations that gene expression changes rapidly in the first hour after dawn (Balcerowicz *et al*., 2021). The first day of sample collection started 24 hours after ZT0 of the first LD (2^nd^ day post-transition). The second day started 5 days later (7^th^ day post-transition). The third started another 5 days later (12^th^ day post-transition). 3 replicates for each time point were taken - with each replicate consisting of the 3rd and 4th true rosette leaves of two separate plants. These were flash frozen in liquid nitrogen and then stored at −80°C. For collections during dark conditions (i.e. ZT16 and ZT20), this work was carried out under green-filtered light.

### RNA-seq and initial data processing

RNA was extracted using the Monarch Total RNA Miniprep Kit (New England Biolabs), according to the manufacturer’s protocol. Residual genomic DNA treatment was removed using the Invitrogen TURBO DNA-free kit (ThermoFisher), using ‘Routine DNase treatment’, according to the manufacturer’s protocol.

Libraries were prepared from high quality RNA by the Genomics Laboratory of the University of York’s Bioscience Technology Facility. Libraries were prepared using the NEBNext Ultra II Directional Library prep kit for Illumina in conjunction with the NEBNext poly(A) magnetic isolation module (New England Biolabs). Library quality was assessed using the Agilent 2100 Bioanalyzer instrument, before pooling at equimolar ratios. Pooled libraries were sent for paired end 150 base sequencing by Novogene (UK) Ltd, on an Illumina NovaSeq 6000.

Before further processing of the raw sequencing data, FastQC v0.11.7 (Andrews, 2010) was used to ensure sample quality passed the expected checks for RNA-seq data. Illumina adapters and low quality bases at the 3’ and 5’ ends were trimmed using CutAdapt v3.4 (Martin, 2011). Reads were quantified using Salmon v1.6.0 (Patro *et al*., 2017) against the TAIR10 transcriptome (Berardini *et al*., 2015).

R (version 4.2.3) was used for analysis of gene expression data (R Core Team, 2023). Transcripts per million (TPM) was used as the measure of relative gene expression across samples. Additionally, TPM levels per transcript isoform were combined to leave only gene-level expression data. 13,156 genes were filtered out before further analysis if all median TPM values (across the three biological replicates per time-of-day and sampling day) were less than or equal to 0.5 (Supplemental Figure 1, Supplemental Table S1). Where log-transformed gene expression is referred to in this text, the transformed expression is log_2_(TPM + 1).

### PCA and GO term overrepresentation

PCA was performed using ‘prcomp’ on log-transformed gene expression on all samples. All GO term overrepresentation was performed using the gprofiler2 R package (version 0.2.2) (Kolberg *et al*., 2020). This called the g:Profiler server (version e111_eg58_p18_f463989d), which utilises the g:SCS multiple testing correction method, and we then applied a significance threshold of 0.05 (Kolberg *et al*., 2023) (Supplemental Tables S2, S3, S6). Related groups of genes were submitted as ‘multi queries’.

### CIBERSORTx

TPM tables were input into the Cell Fraction mode of the CIBERSORTx web interface (https://cibersortx.stanford.edu/, accessed 28 May 2024), using 100 permutations for the permutation test to assess the p-values (Newman *et al*., 2019). The signature matrix came from (Vong *et al*., 2024) and was trained on the single cell RNA-seq data from (Procko *et al*., 2022).

### Rhythmic gene prediction

To predict rhythmic genes, we applied the R package MetaCycle (version 1.2.0) to log-transformed gene expression, only allowing the JTK_cycle method (Hughes *et al*., 2010; Wu *et al*., 2016). Samples at ZT1 were excluded since JTK_cycle required evenly spaced time points. Genes were classified as rhythmic if and only if the ‘BH.Q’ value was less than 0.05. The same method was used to classify the rhythmicity of cell types, where instead we used unscaled cell proportions as inputs.

### Data from previous publications

To identify the putative targets of SNF1 KINASE HOMOLOG10 (KIN10), we took the ‘AGI number’ column from each of the ‘INCREASED’ and ‘DECREASED’ sections from Supplementary Table S3 of ref. (Baena-González *et al*., 2007). Putative targets of CIRCADIAN CLOCK ASSOCIATED1 (CCA1) were collected from Dataset S2 of ref. (Nagel *et al*., 2015). Putative targets of EARLY FLOWERING3 (ELF3) were collected from Supplementary Table 6 of ref. (Ezer *et al*., 2017). Housekeeping genes suggested for developmental and diurnal time series were collected from Supplemental Table 1 of ref. (Czechowski *et al*., 2005). Only gene names which consistent with our processed RNA-seq were used.

## Results

### Gene expression varies more across development than across a diurnal cycle

Since the clock is tightly linked to changes in gene expression over plant development, we first sought to determine whether diurnal or developmental time had a greater impact on gene expression. Principal component analysis (PCA) shows that the first principal component primarily separates out samples based on developmental time, suggesting that variation in gene expression across development dominates variation from diurnal oscillations (Figure 1(a)). As expected, the central day (7^th^ day after transition) lies in between the early (2^nd^) and late (12^th^) days. This agrees with clustering based on correlation between samples (Supplemental Figure 2). 52.5% of variance within the filtered dataset can be explained by the developmental-related principal component – indicating a large transcriptomic scale shift in the 12 days after the transition (Figure 1(a), Supplemental Figure 3).

**Figure 1.**
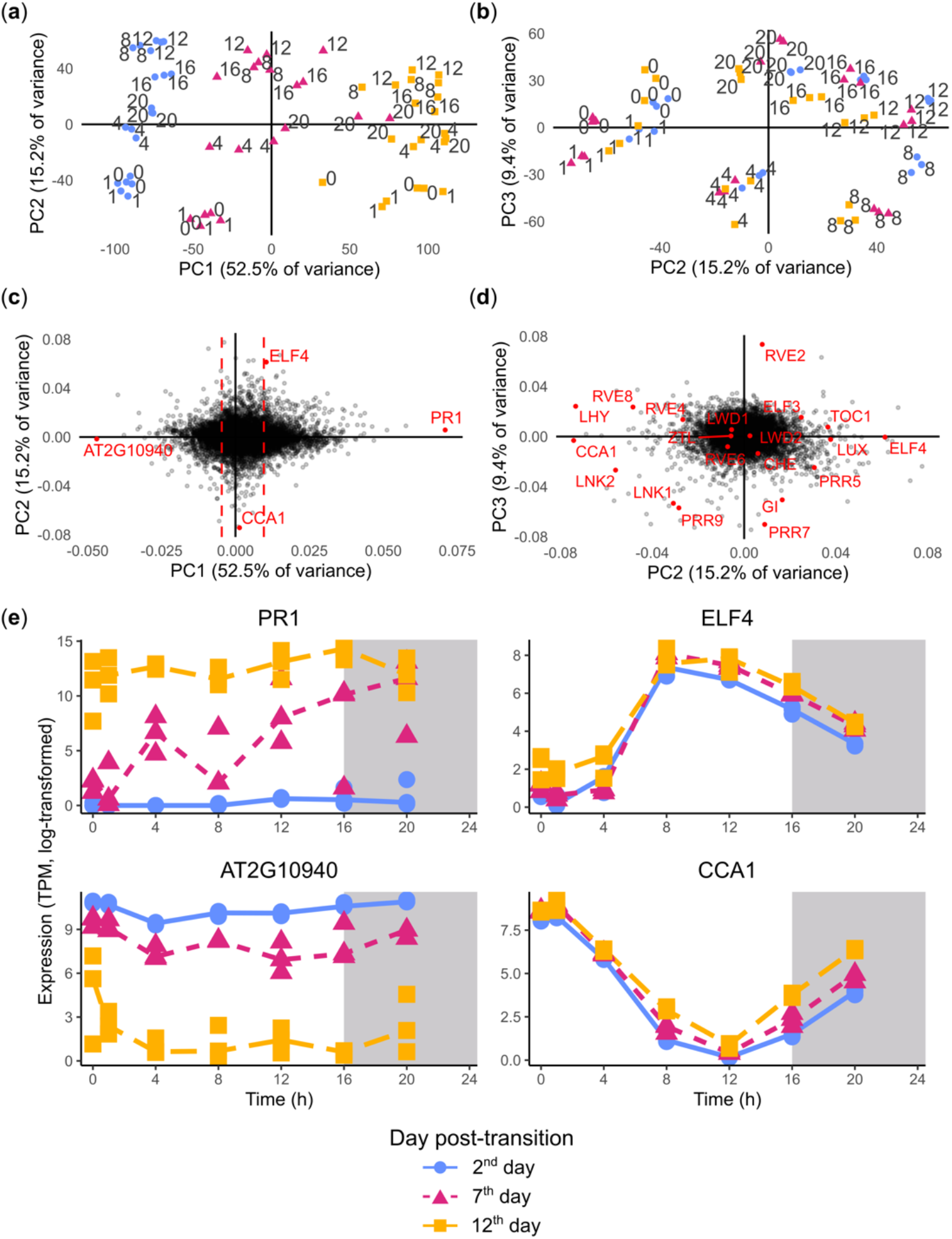
**(a)** Every individual replicate plotted on the first two principal components (PCs), using log-transformed gene expression as inputs. The labelled numbers represent the time of day relative to dawn (zeitgeber time; ZT). Colour and shape represent the day of sample collection, according to the legend at the bottom. **(b)** The same as (a) but for the second and third PCs. **(c)** The distribution of genes according to the loadings on PC1 and PC2. Dashed vertical lines represent the 10% and 90% percentiles on PC1. Genes highlighted in red are plotted in (e). **(d)** The distribution of es according to the loadings on PC2 and PC3. Core clock genes (as defined by Wang et al. 2024) are highlighted in red. **(e)** Expression of the genes with most positive and negative loadings on PC1 and two core clock genes (CCA1 and ELF4). The x-axis represents ZT on each day. Points represent individual biological replicates. Lines are drawn through the median value of the biological replicates.

In the next two principal components (PC2 and PC3), samples clustered based on time-of-day, not day of sampling, with samples forming a circle ordered by the time of day that samples were collected (Figure 1(b)). These results suggest that PCA successfully isolates the developmental and diurnal components of gene expression in our samples.

By analysing the relative contribution (i.e. loadings) of each gene to the first three principal components, we can identify genes that may be associated with developmental (PC1) or diurnal (PC2/3) processes (Figure 1(c), 1(d)). For instance, we can observe that the primary circadian clock genes, as defined by (Wang *et al*., 2024), are correctly ordered in the loadings in Figure 1(d). This includes a precise counter-clockwise ordering of the *PSEUDO-RESPONSE REGULATOR* (*PRR*) genes by the order of their expected phases (Webb *et al*., 2019). The gene with the most positive loading on PC1 was *PATHOGENESIS-RELATED GENE 1* (*PR1*; AT2G14610), a salicylic acid responsive gene whose expression is tightly linked to leaf senescence (Zhang *et al*., 2013). The gene with the most negative loading on PC1 was a proline-rich, cell wall protein AT2G10940 (Duruflé *et al*., 2017). As expected, the expression of *PR1* increases post-transition and the expression of AT2G10940 decrease post-transition (Figure 1(e)). We identified the ‘core clock’ genes, related to the vegetative-to-reproductive transition as defined in (Wang *et al*., 2024), with the most positive and negative loadings on PC2. The clock gene with most positive PC2 loading was *EARLY FLOWERING 4* (*ELF4*; AT2G40080), a member of the evening complex (Nusinow *et al*., 2011; Herrero *et al*., 2012; Ezer *et al*., 2017), which peaked around ZT8 in all days. The clock gene with most negative PC2 loading was *CIRCADIAN CLOCK ASSOCIATED1* (*CCA1*; AT2G46830), which peaked in the early morning in all days. Interestingly, the transcript level of *CCA1* did not appear to decrease across developmental time (i.e. as the leaves aged), contrary to previous reports using qRT-PCR (Song *et al*., 2018). These results suggest that we can use the PCA loadings to identify genes that are primarily developmental or diurnally regulated.

### Developmental changes in gene expression are linked to metabolism, senescence, and kinase-related processes

To determine the broad shifts in gene expression that occur across the developmental timescale, we created two groups of the most early- and late-associated genes. These consisted of genes with the most negative 10% of PC1 loadings (associated with high expression on the 2^nd^ day of sampling) and the most positive 10% (associated with high expression on the 12^th^ day of sampling). GO term overrepresented terms among early-associated genes include ‘photosynthesis’ (GO:0015979) and ‘generation of precursor metabolites and energy’ (GO:0006091), while those overrepresented among late-associated genes include ‘defence response’ (GO:0006952) and ‘plant organ senescence’ (GO:0090693). These are consistent with the vegetative-to-reproductive transition associated GO terms identified in (Hinckley & Brusslan, 2020; Redmond *et al*., 2024) (Supplemental Table S2).

Kinases have been associated with regulating the floral transition, as well as diurnally phosphorylating proteins, forming a bridge between developmental and diurnal processes (Tsai & Gazzarrini, 2012; Uhrig *et al*., 2021). Since ‘protein serine/threonine kinase activity’ (GO:0004674) was overrepresented in the top 10% of genes by positive PC1 loading, we visualised the distribution of all genes related to this GO term (Supplemental Table S2, Supplemental Figure 4(a)). The PC1 (developmental time associated) loadings of these genes were significantly different to 0 (two-sided t-test, t = 17.778, p-value < 2.2e-16) and the mean value was 4.64e-4 (Supplemental Figure 4(a)). However, there was a much smaller deviation from zero for the two diurnal time associated PCs (PC2 and PC3). The loadings on PC2 were not significantly different from 0 (two-sided t-test, t = −1.8644, p-value = 0.06272; mean value −4.47e-4) and the loadings for PC3 were significantly different (two-sided t-test, t = 7.4955, p-value = 2.13e-13; mean value 1.83e-3), but with a much lower effect size than PC1. This suggests that expression of kinases primarily varies over the developmental timescale.

The 4 genes from this category with the highest PC1 loading came from two kinase families. Two of these were *WALL-ASSOCIATED KINASES* (*WAK1, WAK3*), which act as pectin receptors and have roles in defence response and cell expansion (Anderson *et al*., 2001; Kohorn & Kohorn, 2012). A further two were *CYSTEINE-RICH KINASES* (*CRK4, CRK45*). Overexpression of *CRK4* is known to enhance pattern-triggered immunity (Yeh *et al*., 2015). Each of these genes had the property that the expression on day 2 is consistently low throughout the day and the expression of day 12 is consistently high. However, the expression on day 7 spans the range of expression on day 2 and day 12, matching day 2 expression at ZT0 and day 12 expression at ZT20. (Supplemental Figure 4(b)). This could explain the significantly non-zero PC3 loadings of kinases, since PC3 may be incorporating some developmental variation as well as diurnal variation.

### Targets of *SNF1 KINASE HOMOLOG 10 (KIN10)* Show Diurnal and Developmental Expression Changes

Next, we chose to analyse the diurnal and developmental variation of a well-known kinase-containing protein complex that functions as an energy sensor in *Arabidopsis*: the Sucrose non-fermenting-1-related protein kinase (SnRK1) complex (Polge & Thomas, 2007; Broeckx *et al*., 2016), which can entrain the clock in response to sugars (Frank *et al*., 2018) and can regulate growth and development in response to sugar status (Chen *et al*., 2017). The SnRK1 complex consists of three subunits, each of which are encoded from different homologues (Broeckx *et al*., 2016). *SNF1 KINASE HOMOLOG10* (*KIN10*) and *SNF1 KINASE HOMOLOG11 (KIN11*), are protein serine/threonine kinases, which encode the α subunit of SnRK1. Importantly, homologues of all three subunits may be expressed specifically in different developmental stages and environmental conditions (Broeckx *et al*., 2016). Both *KIN10* and *KIN11* had increased expression over the developmental timescale (Figure 2(a)). However, especially in the 12^th^ day, *KIN10* expression had larger diurnal variation than *KIN11*. As for the β subunit, we found that *KIN*β*1* expression had much larger variation over both the diurnal and developmental timescales than expression of its homologues *KIN*β*2* and *KIN*β*3* (Supplemental Figure 5(a)). Finally, the essential gene *KIN*βγ (*KINbeta-gamma, HOMOLOG OF YEAST SUCROSE NONFERMENTING 4, SNF4*) was far more stably expressed over development than the related *KIN*γ (*KINgamma*) (Supplemental Figure 5(b)) (Ramon *et al*., 2013). This could be explained by the fact *KIN*γ is not suspected to form a functional part of the SnRK1 complex, as opposed to its orthologue in mammalian systems (Emanuelle *et al*., 2015).

**Figure 2.**
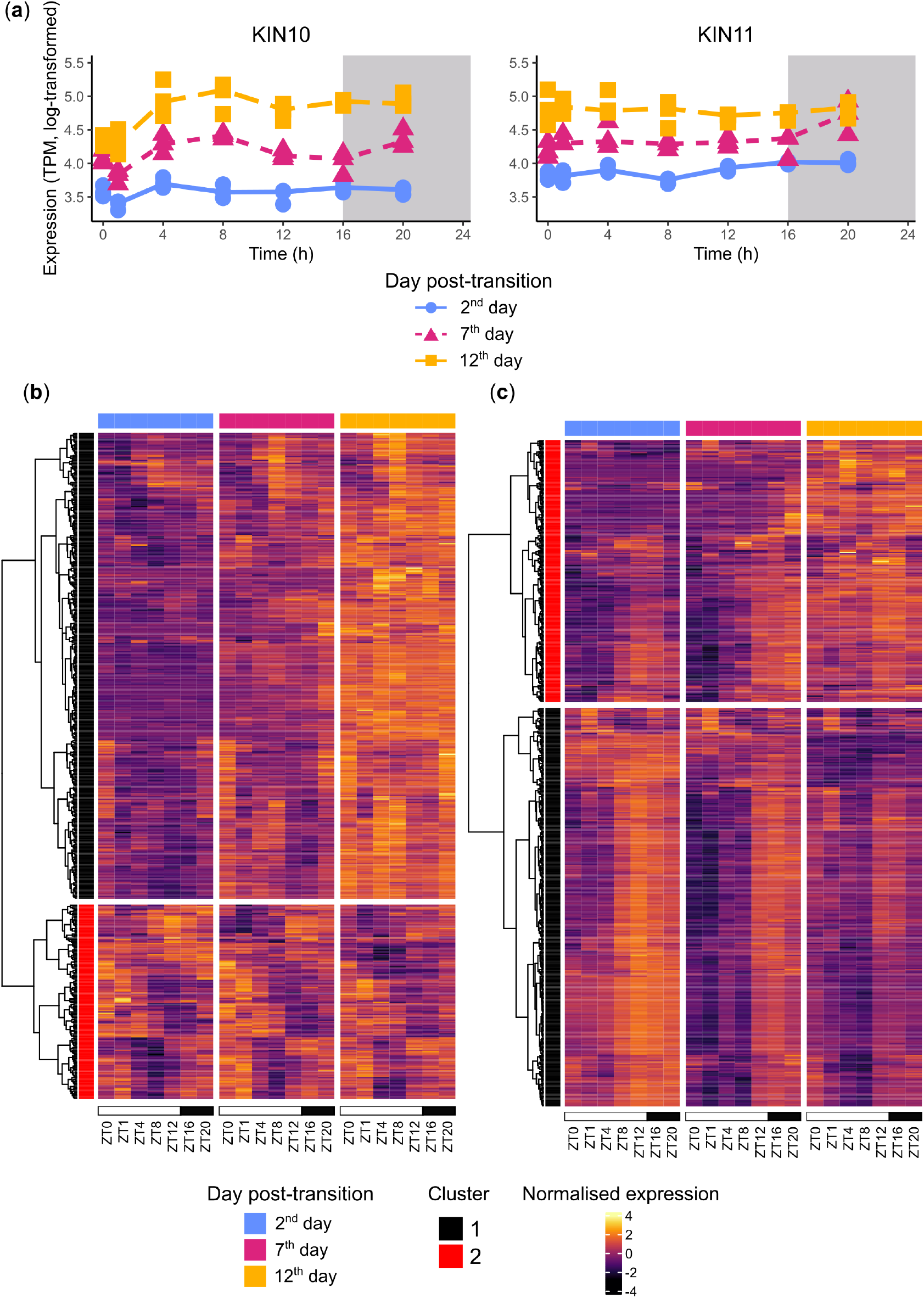
**(a)** Expression of the two genes which encode the *Arabidopsis* orthologue of the α subunit of SnRK1, on the same y-axis so they can be compared. **(b)** Clustering of genes which showed increased expression, in KIN10 overexpressed *Arabidopsis* protoplasts, based on normalised gene expression over both timescales. The method ‘ward.D2’ was used to create the hierarchical clustering of the rows. **(c)** Same as (b) based on genes which showed decreased expression.

Since *KIN10* is known to activate and repress a wide range of pathways (Baena-González *et al*., 2007), we asked how expression of processes that act downstream of *KIN10* changed over diurnal and developmental timescales. We selected genes which increased or decreased expression by transient *KIN10* overexpression in protoplasts and clustered them based on expression over both timescales (Figure 2(b), 2(c)) (Baena-González *et al*., 2007). Both sets of targets clustered into two large subgroups – characterised by their expression patterns over the developmental timescale. In terms of genes with increased expression in the *KIN10* overexpressed lines, cluster 1 showed a large increase over age, which agrees with greater *KIN10* expression over age (Figure 2(a), 2(b)). GO terms significantly overrepresented in this cluster (and not in cluster 2) included ‘response to hypoxia’ (GO:0001666), ‘response to abscisic acid’ (GO:0009737), and ‘carbohydrate metabolic process’ (GO:0005975) (Supplemental Table S3). These represented the range of stress responses *KIN10* is known to affect as well as its control over metabolism (Baena-González *et al*., 2007). Cluster 2 broadly showed a less dramatic increase in expression over age.

In terms of genes with decreased expression in the *KIN10* overexpressed lines, the two clusters showed strikingly different expression. Cluster 1 expression broadly decreased in expression over age, which agrees with increasing *KIN10* expression (Figure 2(a)). The majority of these targets also peaked at ZT12 or later in the day, when starch is degraded to provide a carbohydrate source (Figure 2(c)) (Streb & Zeeman, 2012). This cluster appeared to represent the sink-to-source transition of ageing leaves, since both ‘primary metabolic process’ (GO:0044238) and ‘cellular nitrogen compound metabolic process’ (GO:0034641) were significantly overrepresented (Supplemental Table S3) (Havé *et al*., 2017). Contrarily, cluster 2 showed a broad increase in expression. This suggests that *KIN10* repression in protoplasts does not always predict repression in ageing plants, at least at the transcriptional level. Our results suggest that SnRK1 sub-components primarily changed their transcriptional profiles developmentally, but that most genes that were downregulated in response to *KIN10* expression have consistent diurnal expression patterns.

### Many targets of core clock genes vary their expression patterns over development

Although PCs 2 and 3 (the diurnal components) accounted for only 24.6% of the variation in our dataset and the diurnal variation of *KIN10* expression was relatively small, metabolic signals are known to be both inputs and outputs of the circadian clock (Shin *et al*., 2017; Frank *et al*., 2018; Cervela-Cardona *et al*., 2021). Additionally, senescence-associated changes in period and phase have been observed in both the core clock genes and the wider circadian transcriptome (Kim *et al*., 2016, 2018; Buckley *et al*., 2024). We noticed that most clock genes had expression patterns that were consistent across development, under our fixed long-day conditions (Supplemental Figure 6). This prompts the questions of whether clock gene *targets* had stable expression over development and whether they were rhythmic in each day after the SD to LD transition.

*CIRCADIAN-CLOCK ASSOCIATED 1* (*CCA1*) is a core clock gene whose expression peaks in the early morning under long days (Figure 3(a)) (Webb *et al*., 2019; Wang *et al*., 2024). We collected predicted *CCA1* targets from a previous ChIP-seq study, where seedlings were grown under 12h light/ 12h dark conditions (Nagel *et al*., 2015). We observed that the median expression of *CCA1*, at every time of day, increased slightly over development (Figure 3(a)). Surprisingly, we found that clusters targets of *CCA1* demonstrated increasing, decreasing, and stable mean expression per day of the experiment (Figure 3(b), 3(c), Supplemental Table S5). Further, we found that some clusters of targets became less rhythmic over age (e.g. clusters B and E), according to JTK_cycle, while some clusters maintained high levels (>80%) of rhythmicity (cluster C) (Hughes *et al*., 2010).

**Figure 3.**
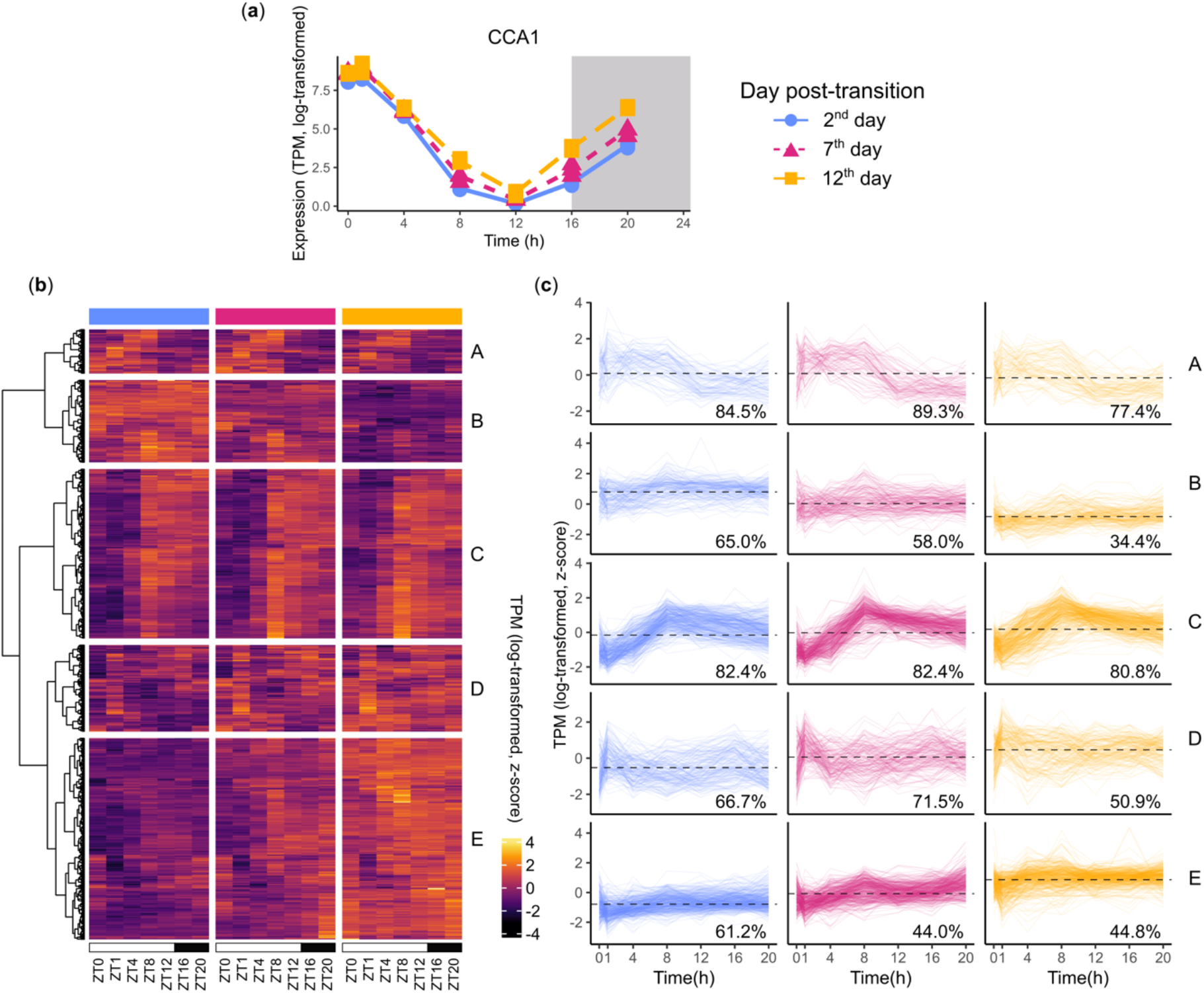
**(a)** Expression of CCA1. **(b)** Heatmap of targets from CCA1, from ChIP-seq data in (Nagel et al., 2015). Values represent log-transformed then z-scored gene expression, so expression can be compared across different genes. **(c)** Timeseries of targets, grouped by cluster. These follow the same vertical order as in (b). Dashed lines represent mean expression over all genes plotted and all timepoints. The percentages refer to the proportion of genes which were significantly rhythmic in each day, as predicted by JTK_cycle.

To show if these complicated patterns of regulation were preserved across other clock genes, we looked at predicted targets of another clock gene, *ELF3*, from (Ezer *et al*., 2017). *ELF3* is a component of the EC and its expression peaks around ZT8 under long days, 8 hours later than *CCA1* (Figure 4(a)) (Ezer *et al*., 2017; Webb *et al*., 2019). Intriguingly, as with *CCA1*, many clusters showed a low proportion of rhythmic genes (<51%) on the 12^th^ day after the SD to LD transition. As for highly rhythmic clusters, they showed varying behaviour over development. Clusters C and D were both highly rhythmic on the 2^nd^, 7^th^, and 12^th^ days and their expression broadly peaked before ZT8 on each day, suggesting that they were repressed by the EC (Figure 4(b), 4(c); Supplemental Table S4, S5). The mean expression of cluster C over each day slightly increased whereas this decreased for cluster D. Both clusters had an overrepresentation of ‘photosystem I’ (GO:0009522) and multiple other photosynthesis related terms (Supplemental Table S6). However, many regulatory GO terms were overrepresented in cluster C but not in cluster D, such as ‘regulation of biosynthetic process’ (GO:0009889) and ‘response to red or far red light’ (GO:0009639). *PHYTOCHROME-INTERACTING FACTOR5* (*PIF5*), which is phosphorylated and degraded in response to red-light, was in cluster C (Shen *et al*., 2007). This suggests that the interplay between the EC and *PHYTOCHROME B* (*PHYB*) (Ezer *et al*., 2017) may be stable over development.

**Figure 4.**
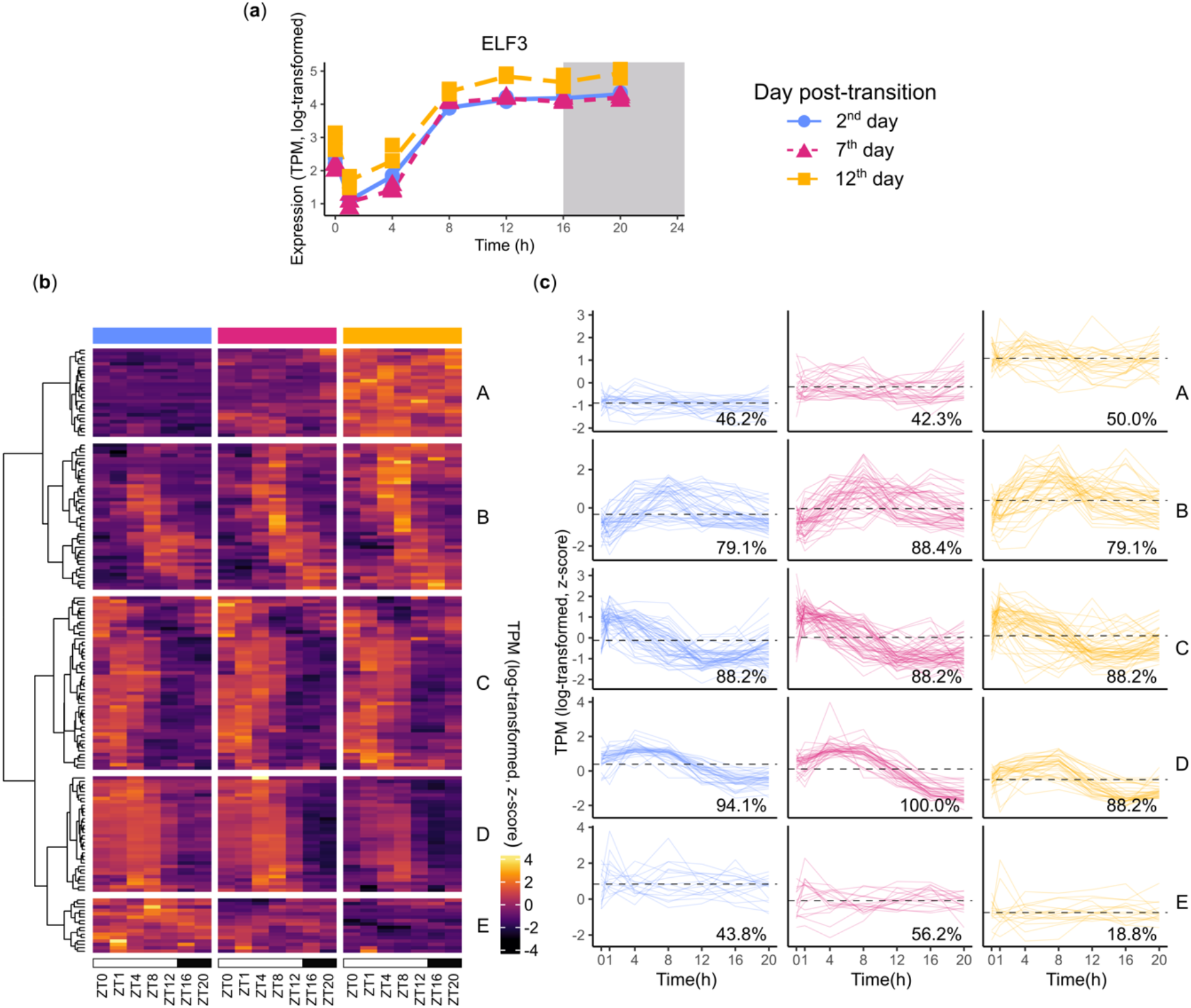
**(a)** Expression of ELF3. **(b)** Heatmap of targets from ELF3, from ChIP-seq data in (Ezer et al., 2017). Values represent log-transformed then z-scored gene expression, so expression can be compared across different genes. **(c)** Timeseries of targets, grouped by cluster, with the same labels as in Figure 3(c).

**Figure 5.**
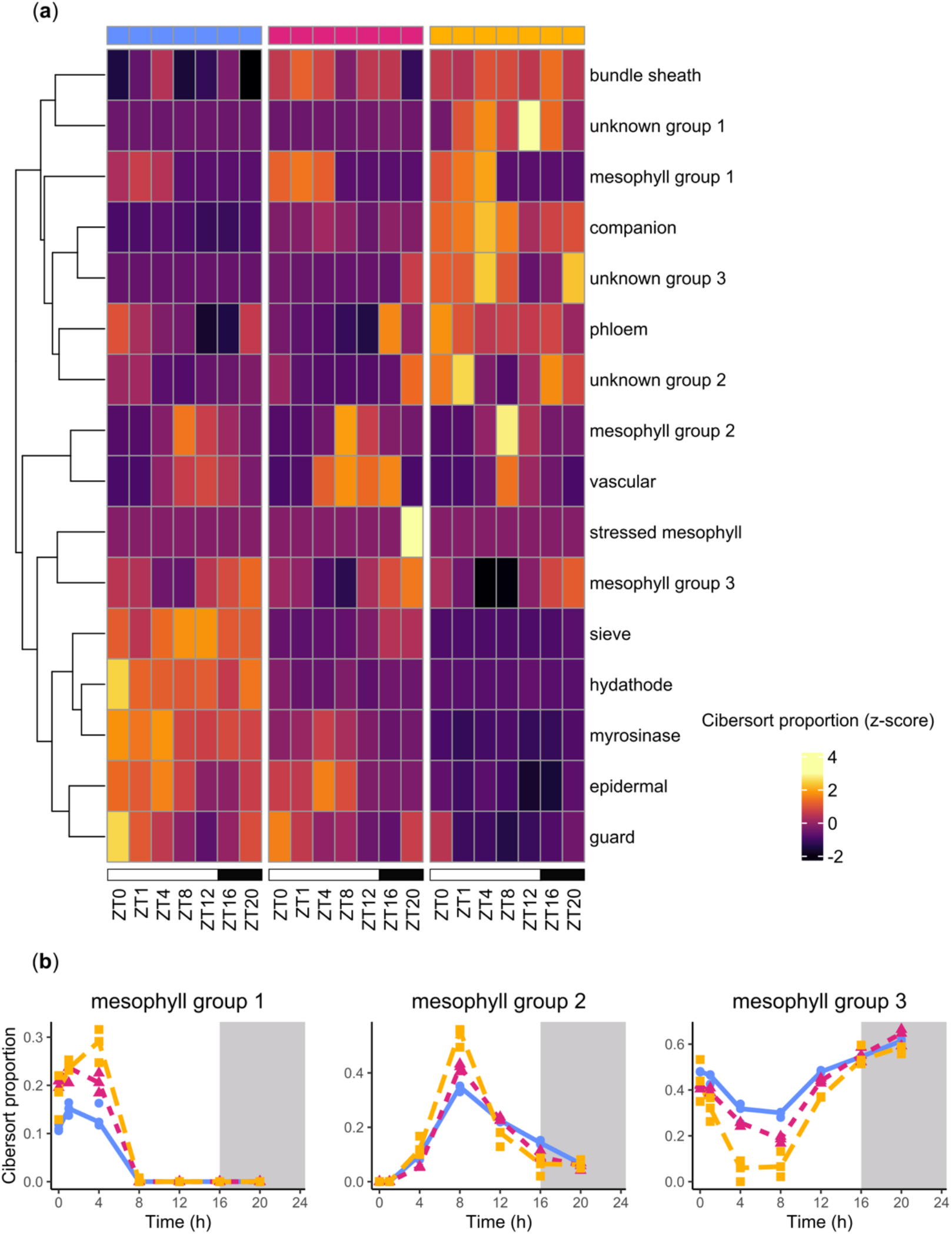
**(a)** Heatmap of cibersort outputs. Values represent the predicted proportion of gene expression from each sample that came from each cell type. Values show represent the median value of the three biological replicates, which have been scaled and then values higher than 3 or lower than −3 have been truncated. Rows have been clustered with hierarchical clustering using the “complete” method. **(b)** The proportion of three different groups of mesophyll cell types. Points represent the values for the individual replicates and lines are drawn through median values.

### Cell-type specific activity changes over diurnal and developmental timescales

Since different clock components are enriched in various *Arabidopsis* leaf cell types (Nohales, 2021; Davis *et al*., 2022), we wanted to know how cell-specific diurnal expression of genes changes over age. Our previous work adapted a method, CIBERSORTx, to estimate cell type-specific activity in bulk leaf RNA-seq samples (Newman *et al*., 2019; Vong *et al*., 2024). Most cell types are predicted to have rhythmic activity on all three days of sampling (JTK_cycle, BH.q value < 0.05; Supplemental Figure 7, Supplemental Table S8). However, some cell types with low activity early in development are predicted not to be rhythmic, then become rhythmic later in development, such as ‘Unknown Group 3’. Vong *et al*., 2024 suggested that these groups may represent stressed or senescent cell types. Our results also confirm that phloem and the unknown cell groups were expressed late in development, while the epidermal cells were primarily transcriptionally active early in development (Figure 4(a)). We also confirm consistent diurnal oscillations in guard cell and vascular cell transcriptional activity.

Coupling between the circadian clocks in mesophyll and vasculature has been demonstrated (Endo *et al*., 2014). We therefore sought to understand the behaviour of subgroups of mesophyll. For instance, there were three groups of mesophyll cells that were each primarily active at different times of day and different stages of development. Our results demonstrate that the phase of their expression is consistent throughout development, but that the peak expression increases in groups 1 and 2 over development and the minimum decreases in group 3 (Figure 4(b)). These results highlight that any differentially expressed genes over diurnal or developmental time may be a consequence of oscillations in which cells are transcriptionally active, rather than a homogenous change of expression across all cell types.

### Known housekeeping genes display variation over both developmental and diurnal timescales

Our previous analysis found many examples of genes that change their expression diurnally and / or over development. However, it is also important to find sets of genes that are stably expressed under both conditions. These are important controls in experiments like reverse transcription followed by quantitative polymerase chain reaction (qRT-PCR), where gene expression must be normalised to a housekeeper gene (Czechowski *et al*., 2005; Souček *et al*., 2017). Researchers often refer to well-known lists of housekeeping genes, which are suggested to be stable under relevant experimental conditions. We calculated the coefficient of variation (CV; standard deviation divided by mean) from our dataset of the suggested housekeeping genes for developmental and diurnal experiments from ref. (Czechowski *et al*., 2005). A high CV value indicates that a gene shows high biological variability gene expression is not comparable across different times of day or developmental stages. Although the mean CV value of the housekeeping genes was lower than the mean CV value across all genes, many housekeeping genes still showed undesirable levels of variation (Figure 6(a)). For example, two commonly used housekeeping genes, *PROTEIN PHOSPHATASE 2A* (*PP2A) SUBUNIT3* and *YELLOW-LEAF-SPECIFIC GENE8 (YLS8)*, had large expression changes over development (Figure 6(b)). *YLS8* approximately doubled in expression from the 2^nd^ day of sampling to the 12^th^ day. However, some housekeeping genes did show promisingly low CV scores. Genes *ARABIDOPSIS RNA POLYMERASE B13*.*6* (*ATRPB13*.*6;* AT3G52090), AT3G03070, and AT3G12260 had the lowest CV values across all housekeeping genes (Figure 6(a)). Despite the low CV values, we observed that these genes still displayed variation across the diurnal timescale, when visualised on a relative expression scale (Figure 6(c)). We suggest that combinations of these genes may be useful as controls for qRT-PCR.

**Figure 6.**
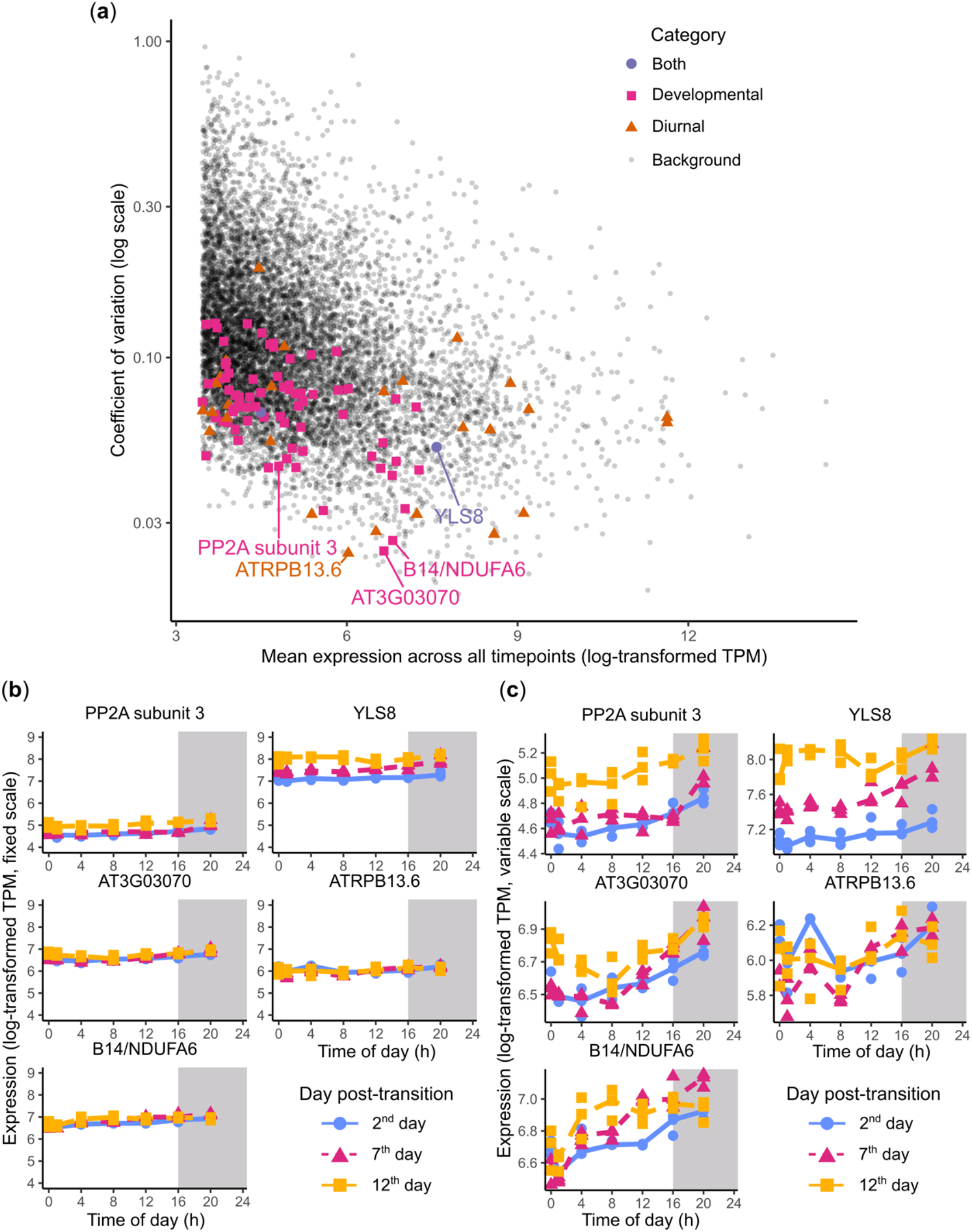
**(a)** Coefficient of variation (CV; calculated by standard deviation / mean) versus mean expression of genes. Both values are calculated including all 63 samples (all replicates at all timepoints). Only genes with a TPM > 10 are shown. Samples which were suggested by (Czechowski et al., 2005) as good housekeeping qPCR genes for diurnal experiments or developmental experiments (or both) are highlighted. **(b)** Expression of 5 housekeeping genes selected from (a). PP2A subunit 3 and YLS8 have relatively high CV values, and the other genes have the lowest 3 CV values. The y axes are fixed so the developmental variation can be compared across genes. **(c)** Same as (b), except the y axes vary to show diurnal variation.

## Discussion

We measured gene expression via RNA-seq in ageing plants, across both diurnal and developmental timescales. Using PCA, we observed that most of the variation in samples can be explained by a developmental PC, with a smaller contribution from two diurnal PCs (Figure 1). These PCA loadings help us establish the extent to which any individual gene varies its expression across diurnal and developmental timescales.

Although kinases are thought to associate diurnal and developmental programmes, we found that they primarily change their expression across developmental scales. However, many genes whose expression changes are perturbed by *KIN10* overexpression have diurnal expression patterns, which is consistent with previous suggestions that there is crosstalk between the SnRK1 complex and circadian processes (Shin *et al*., 2017; Chen *et al*., 2017; Frank *et al*., 2018). This highlights that circadian control of primary metabolic processes, including turnover of carbon and nitrogen reserves, remains important during the onset of senescence (Flis *et al*., 2019). Due to the fixed photoperiod that we used throughout the experiment, we could not observe photoperiod-dependent effects, which have been shown to affect the diurnal timing of metabolic processes (Flis *et al*., 2016; Alexandre Moraes *et al*., 2022). Future work could test whether these effects are consistent across different stages of plant development.

We did not observe changes in the phase or period of core clock genes (Supplemental Figure 6). This is unsurprising given that our ageing plants were exposed to LD (entrainment) conditions on the days of sampling, rather than free-running conditions, and it highlights the stability of the central oscillator. However, it has previously been observed that the clock can have different free-running periods in younger versus older leaves (Kim *et al*., 2016). This is consistent with our and previous reported observations that metabolic signalling inputs to the clock, such as *KIN10*, have different levels of expression across plant development (Baena-González *et al*., 2007; Frank *et al*., 2018).

Despite the relatively stable expression of *CCA1* and *ELF3* over development, their predicted targets cluster into groups that either decrease or increase in mean expression (Figure 3(b), Figure 4(b)). This could be because of changing patterns of core clock gene binding to regulatory regions of the genome. However, these patterns may instead result from downstream processing of circadian signals. For example, the clock influences senescence through both transcriptional and *miR164*-mediated post-translation control of *ORESARA1* (*ORE1*) (Kim *et al*., 2018). The extent of these post-translations mechanisms could be assessed by measuring the changes in the diurnal proteome and phosphoproteome in ageing plants (Uhrig *et al*., 2021). Our work suggests that there is value in spanning both developmental and diurnal timescales and suggests that other complementary datasets, such as ChIP-seq, proteomics, and phosphoproteomics, should be performed at more than one timepoint per day and multiple stages of leaf development (Nagel *et al*., 2015; O’Malley *et al*., 2016; Ezer *et al*., 2017; Li *et al*., 2023).

Diurnal rhythms of gene expression have previously been observed in different cell types, through luciferase assays, qRT-PCR, and single-nucleus RNA-seq (Endo *et al*., 2014; Greenwood *et al*., 2022; Qin *et al*., 2023). Here, we utilised CIBERSORTx to deconvolute gene expression from bulk RNA-seq data into contributions from leaf-specific cell types, allowing us to see how diurnal patterns change over development (Newman *et al*., 2019). We identified some cell types that became rhythmic only at the latest stages of development. We also showed that subgroups of mesophyll cells exhibit different timings of expression across both timescales. It remains to be seen whether the reported coordination of the clock across cell types is maintained across development (Endo *et al*., 2014; Gould *et al*., 2018; Schmal *et al*., 2019).

We provide a valuable resource for the community to search for appropriate housekeeping genes when researching the clock in ageing plants. Although existing housekeeping genes show lower variability than background gene expression, we recommend that researchers use at least one of our three suggested genes. Our suggested genes showed little variation across the developmental timescale. Using multiple housekeeping genes to normalise qRT-PCR data may counterbalance the unavoidable diurnal variation of housekeeping genes (Vandesompele *et al*., 2002).

## Acknowledgements

This project was undertaken on the Viking Cluster, which is a high-performance compute facility provided by the University of York. The authors are grateful for computational support from the University of York High Performance Computing service, Viking and the Research Computing team. They also acknowledge support from the Horticulture Facility and the Genomics Laboratory in the Technology Facility at the Department of Biology, University of York.

## Funding

This work was supported by funding from the Biotechnology and Biological Sciences Research Council (BBSRC)—D.E. and S.J.D.: BB/V006665/1; D.E.: BB/S506795/1; and S.J.D.: BB/N018540/1. The authors also acknowledge BBSRC White Rose DTP studentships (BB/M011151/1 and BB/T007222/1) to J.R. (Ref.: 1792522) and E.J.R. (Ref.: 2444228). D.E. was also supported by the Royal Society (RGS\R2\212345).

## Competing interests

No competing interests are declared.

## Author contributions

E.J.R.: conceptualization, methodology, software, formal analysis, investigation, writing— original draft, and visualization. J.R.: conceptualization, investigation, and writing—review and editing. S.J.D.: writing—review and editing, supervision, and funding acquisition. D.E.: conceptualization, methodology, writing—original draft, writing—review and editing, supervision, and funding acquisition.

## Data availability

Raw and processed sequencing data have been deposited in the NCBI Gene Expression Omnibus (GEO) database under accession number GSE242964 (https://www.ncbi.nlm.nih.gov/geo/query/acc.cgi?acc=GSE242964). Scripts used to produce the figures have been deposited in a GitHub repository https://github.com/stressedplants/AgeingCircadian

## Notes

https://www.ncbi.nlm.nih.gov/geo/query/acc.cgi?acc=GSE242964

### Competing Interest Statement

The authors have declared no competing interest.

### Summary of Updates

Additional key references have been added and the discussion has been expanded. Certain textual elements elsewhere have been edited for clarity.

## References

Alexandre Moraes T, Mengin V, Peixoto B, Encke B, Krohn N, Höhne M, Krause U, Stitt M. 2022. The circadian clock mutant lhy cca1 elf3 paces starch mobilization to dawn despite severely disrupted circadian clock function. Plant physiology 189: 2332–2356.

Anderson CM, Wagner TA, Perret M, He ZH, He D, Kohorn BD. 2001. WAKs: cell wall-associated kinases linking the cytoplasm to the extracellular matrix. Plant molecular biology 47: 197–206.

Andrade L, Lu Y, Cordeiro A, Costa JMF, Wigge PA, Saibo NJM, Jaeger KE. 2022. The evening complex integrates photoperiod signals to control flowering in rice. Proceedings of the National Academy of Sciences of the United States of America 119: e2122582119.

Andrews S. 2010. FastQC: a quality control tool for high throughput sequence data.

Anwer MU, Davis A, Davis SJ, Quint M. 2020. Photoperiod sensing of the circadian clock is controlled by EARLY FLOWERING 3 and GIGANTEA. The Plant journal: for cell and molecular biology 101: 1397–1410.

Atamian HS, Creux NM, Brown EA, Garner AG, Blackman BK, Harmer SL. 2016. Circadian regulation of sunflower heliotropism, floral orientation, and pollinator visits. Science 353: 587–590.

Baena-González E, Rolland F, Thevelein JM, Sheen J. 2007. A central integrator of transcription networks in plant stress and energy signalling. Nature 448: 938–942.

Balcerowicz M, Mahjoub M, Nguyen D, Lan H, Stoeckle D, Conde S, Jaeger KE, Wigge PA, Ezer D. 2021. An early-morning gene network controlled by phytochromes and cryptochromes regulates photomorphogenesis pathways in Arabidopsis. Molecular plant 14: 983–996.

Berardini TZ, Reiser L, Li D, Mezheritsky Y, Muller R, Strait E, Huala E. 2015. The Arabidopsis information resource: Making and mining the “gold standard” annotated reference plant genome. Genesis 53: 474–485.

Broeckx T, Hulsmans S, Rolland F. 2016. The plant energy sensor: evolutionary conservation and divergence of SnRK1 structure, regulation, and function. Journal of experimental botany 67: 6215–6252.

Buckley CR, Boyte JM, Albiston RL, Hyles J, Beasley JT, Johnson AAT, Trevaskis B, Fournier-Level A, Haydon MJ. 2024. A circadian transcriptional sub-network and EARLY FLOWERING 3 control timing of senescence and grain nutrition in bread wheat. bioRxiv: 2024.02.19.580927.

Cervela-Cardona L, Yoshida T, Zhang Y, Okada M, Fernie A, Mas P. 2021. Circadian Control of Metabolism by the Clock Component TOC1. Frontiers in plant science 12: 683516.

Chen L, Su Z-Z, Huang L, Xia F-N, Qi H, Xie L-J, Xiao S, Chen Q-F. 2017. The AMP-activated protein kinase KIN10 is involved in the regulation of autophagy in Arabidopsis. Frontiers in plant science 8: 1201.

Corbesier L, Gadisseur I, Silvestre G, Jacqmard A, Bernier G. 1996. Design in Arabidopsis thaliana of a synchronous system of floral induction by one long day. The Plant journal: for cell and molecular biology 9: 947–952.

Covington MF, Maloof JN, Straume M, Kay SA, Harmer SL. 2008. Global transcriptome analysis reveals circadian regulation of key pathways in plant growth and development. Genome biology 9: R130.

Czechowski T, Stitt M, Altmann T, Udvardi MK, Scheible W-R. 2005. Genome-wide identification and testing of superior reference genes for transcript normalization in Arabidopsis. Plant physiology 139: 5–17.

Davis W, Endo M, Locke JCW. 2022. Spatially specific mechanisms and functions of the plant circadian clock. Plant physiology 190: 938–951.

Dodd AN, Salathia N, Hall A, Kévei E, Tóth R, Nagy F, Hibberd JM, Millar AJ, Webb AAR. 2005. Plant circadian clocks increase photosynthesis, growth, survival, and competitive advantage. Science 309: 630–633.

Duruflé H, Hervé V, Balliau T, Zivy M, Dunand C, Jamet E. 2017. Proline Hydroxylation in Cell Wall Proteins: Is It Yet Possible to Define Rules? Frontiers in plant science 8: 1802.

Emanuelle S, Hossain MI, Moller IE, Pedersen HL, van de Meene AML, Doblin MS, Koay A, Oakhill JS, Scott JW, Willats WGT, et al. 2015. SnRK1 from Arabidopsis thaliana is an atypical AMPK. The Plant journal: for cell and molecular biology 82: 183–192.

Endo M, Shimizu H, Nohales MA, Araki T, Kay SA. 2014. Tissue-specific clocks in Arabidopsis show asymmetric coupling. Nature 515: 419–422.

Ezer D, Jung J-H, Lan H, Biswas S, Gregoire L, Box MS, Charoensawan V, Cortijo S, Lai X, Stöckle D, et al. 2017. The evening complex coordinates environmental and endogenous signals in Arabidopsis. Nature plants 3: 17087.

Flis A, Mengin V, Ivakov AA, Mugford ST, Hubberten H-M, Encke B, Krohn N, Höhne M, Feil R, Hoefgen R, et al. 2019. Multiple circadian clock outputs regulate diel turnover of carbon and nitrogen reserves: Metabolism in five circadian clock mutants. Plant, cell & environment 42: 549–573.

Flis A, Sulpice R, Seaton DD, Ivakov AA, Liput M, Abel C, Millar AJ, Stitt M. 2016. Photoperiod-dependent changes in the phase of core clock transcripts and global transcriptional outputs at dawn and dusk in Arabidopsis. Plant, cell & environment 39: 1955–1981.

Fowler S, Lee K, Onouchi H, Samach A, Richardson K, Morris B, Coupland G, Putterill J. 1999. GIGANTEA: a circadian clock-controlled gene that regulates photoperiodic flowering in Arabidopsis and encodes a protein with several possible membrane-spanning domains. The EMBO journal 18: 4679–4688.

Frank A, Matiolli CC, Viana AJC, Hearn TJ, Kusakina J, Belbin FE, Wells Newman D, Yochikawa A, Cano-Ramirez DL, Chembath A, et al. 2018. Circadian Entrainment in Arabidopsis by the Sugar-Responsive Transcription Factor bZIP63. Current biology: CB 28: 2597–2606.e6.

Gould PD, Domijan M, Greenwood M, Tokuda IT, Rees H, Kozma-Bognar L, Hall AJ, Locke JC. 2018. Coordination of robust single cell rhythms in the Arabidopsis circadian clock via spatial waves of gene expression. eLife 7.

Greenwood M, Hall AJW, Locke JCW. 2022. High Spatial Resolution Luciferase Imaging of the Arabidopsis thaliana Circadian Clock. Methods in molecular biology 2398: 47–55.

Havé M, Marmagne A, Chardon F, Masclaux-Daubresse C. 2017. Nitrogen remobilization during leaf senescence: lessons from Arabidopsis to crops. Journal of experimental botany 68: 2513–2529.

Hayama R, Sarid-Krebs L, Richter R, Fernández V, Jang S, Coupland G. 2017. PSEUDO RESPONSE REGULATORs stabilize CONSTANS protein to promote flowering in response to day length. The EMBO journal 36: 904–918.

Herrero E, Kolmos E, Bujdoso N, Yuan Y, Wang M, Berns MC, Uhlworm H, Coupland G, Saini R, Jaskolski M, et al. 2012. EARLY FLOWERING4 recruitment of EARLY FLOWERING3 in the nucleus sustains the Arabidopsis circadian clock. The Plant cell 24: 428–443.

Hinckley WE, Brusslan JA. 2020. Gene expression changes occurring at bolting time are associated with leaf senescence in Arabidopsis. Plant direct 4: e00279.

Hughes ME, Hogenesch JB, Kornacker K. 2010. JTK_CYCLE: an efficient nonparametric algorithm for detecting rhythmic components in genome-scale data sets. Journal of biological rhythms 25: 372–380.

Inoue K, Araki T, Endo M. 2018. Circadian clock during plant development. Journal of plant research 131: 59–66.

Kim H, Kim HJ, Vu QT, Jung S, McClung CR, Hong S, Nam HG. 2018. Circadian control of ORE1 by PRR9 positively regulates leaf senescence in Arabidopsis. Proceedings of the National Academy of Sciences of the United States of America 115: 8448–8453.

Kim H, Kim Y, Yeom M, Lim J, Nam HG. 2016. Age-associated circadian period changes in Arabidopsis leaves. Journal of experimental botany 67: 2665–2673.

Klingenberg CP. 2019. Phenotypic Plasticity, Developmental Instability, and Robustness: The Concepts and How They Are Connected. Frontiers in Ecology and Evolution 7.

Kohorn BD, Kohorn SL. 2012. The cell wall-associated kinases, WAKs, as pectin receptors. Frontiers in plant science 3: 88.

Kolberg L, Raudvere U, Kuzmin I, Adler P, Vilo J, Peterson H. 2023. g:Profiler-interoperable web service for functional enrichment analysis and gene identifier mapping (2023 update). Nucleic acids research 51: W207–W212.

Kolberg L, Raudvere U, Kuzmin I, Vilo J, Peterson H. 2020. gprofiler2 -- an R package for gene list functional enrichment analysis and namespace conversion toolset g:Profiler. F1000Research 9.

Krzymuski M, Andrés F, Cagnola JI, Jang S, Yanovsky MJ, Coupland G, Casal JJ. 2015. The dynamics of FLOWERING LOCUS T expression encodes long-day information. The Plant journal: for cell and molecular biology 83: 952–961.

Li M, Yao T, Lin W, Hinckley WE, Galli M, Muchero W, Gallavotti A, Chen J-G, Huang S-SC. 2023. Double DAP-seq uncovered synergistic DNA binding of interacting bZIP transcription factors. Nature communications 14: 2600.

Martin M. 2011. Cutadapt removes adapter sequences from high-throughput sequencing reads. EMBnet.journal 17: 10–12.

Nagel DH, Doherty CJ, Pruneda-Paz JL, Schmitz RJ, Ecker JR, Kay SA. 2015. Genome-wide identification of CCA1 targets uncovers an expanded clock network in Arabidopsis. Proceedings of the National Academy of Sciences of the United States of America 112: E4802–10.

Newman AM, Steen CB, Liu CL, Gentles AJ, Chaudhuri AA, Scherer F, Khodadoust MS, Esfahani MS, Luca BA, Steiner D, et al. 2019. Determining cell type abundance and expression from bulk tissues with digital cytometry. Nature biotechnology 37: 773–782.

Nohales MA. 2021. Spatial Organization and Coordination of the Plant Circadian System. Genes 12.

Nozue K, Covington MF, Duek PD, Lorrain S, Fankhauser C, Harmer SL, Maloof JN. 2007. Rhythmic growth explained by coincidence between internal and external cues. Nature 448: 358–361.

Nusinow DA, Helfer A, Hamilton EE, King JJ, Imaizumi T, Schultz TF, Farré EM, Kay SA. 2011. The ELF4-ELF3-LUX complex links the circadian clock to diurnal control of hypocotyl growth. Nature 475: 398–402.

O’Malley RC, Huang S-SC, Song L, Lewsey MG, Bartlett A, Nery JR, Galli M, Gallavotti A, Ecker JR. 2016. Cistrome and Epicistrome Features Shape the Regulatory DNA Landscape. Cell 165: 1280–1292.

Park DH, Somers DE, Kim YS, Choy YH, Lim HK, Soh MS, Kim HJ, Kay SA, Nam HG. 1999. Control of circadian rhythms and photoperiodic flowering by the Arabidopsis GIGANTEA gene. Science (New York, N.Y.) 285: 1579–1582.

Patro R, Duggal G, Love MI, Irizarry RA, Kingsford C. 2017. Salmon provides fast and bias-aware quantification of transcript expression. Nature methods 14: 417–419.

Petersen J, Rredhi A, Szyttenholm J, Mittag M. 2022. Evolution of circadian clocks along the green lineage. Plant physiology 190: 924–937.

Polge C, Thomas M. 2007. SNF1/AMPK/SnRK1 kinases, global regulators at the heart of energy control? Trends in plant science 12: 20–28.

Procko C, Lee T, Borsuk A, Bargmann BOR, Dabi T, Nery JR, Estelle M, Baird L, O’Connor C, Brodersen C, et al. 2022. Leaf cell-specific and single-cell transcriptional profiling reveals a role for the palisade layer in UV light protection. The Plant cell 34: 3261–3279.

Qin Y, Liu Z, Gao S, Long Y, Zhu X, Liu B, Gao Y, Xie Q, Nohales MA, Xu X, et al. 2023. A day in the life of Arabidopsis:24-hour time-lapse single-nucleus transcriptomics reveal cell-type specific circadian rhythms. bioRxiv: 2023.12.09.570919.

R Core Team. 2023. R: A language and environment for statistical computing.

Ramon M, Ruelens P, Li Y, Sheen J, Geuten K, Rolland F. 2013. The hybrid four-CBS-domain KINβγ subunit functions as the canonical γ subunit of the plant energy sensor SnRK1. The Plant journal: for cell and molecular biology 75: 11–25.

Redmond EJ, Ronald J, Davis SJ, Ezer D. 2024. Single-plant-omics reveals the cascade of transcriptional changes during the vegetative-to-reproductive transition. The plant cell: 2023.09.11.557157.

Rees H, Rusholme-Pilcher R, Bailey P, Colmer J, White B, Reynolds C, Ward SJ, Coombes B, Graham CA, de Barros Dantas LL, et al. 2022. Circadian regulation of the transcriptome in a complex polyploid crop. PLoS biology 20: e3001802.

Romanowski A, Schlaen RG, Perez-Santangelo S, Mancini E, Yanovsky MJ. 2020. Global transcriptome analysis reveals circadian control of splicing events in Arabidopsis thaliana. The Plant journal: for cell and molecular biology 103: 889–902.

Rubin MJ, Brock MT, Davis AM, German ZM, Knapp M, Welch SM, Harmer SL, Maloof JN, Davis SJ, Weinig C. 2017. Circadian rhythms vary over the growing season and correlate with fitness components. Molecular ecology 26: 5528–5540.

Sakuraba Y, Jeong J, Kang M-Y, Kim J, Paek N-C, Choi G. 2014. Phytochrome-interacting transcription factors PIF4 and PIF5 induce leaf senescence in Arabidopsis. Nature communications 5: 4636.

Sawa M, Kay SA. 2011. GIGANTEA directly activates Flowering Locus T in Arabidopsis thaliana. Proceedings of the National Academy of Sciences of the United States of America 108: 11698–11703.

Schmal C, Ono D, Myung J, Pett JP, Honma S, Honma K-I, Herzel H, Tokuda IT. 2019. Weak coupling between intracellular feedback loops explains dissociation of clock gene dynamics. PLoS computational biology 15: e1007330.

Seaton DD, Smith RW, Song YH, MacGregor DR, Stewart K, Steel G, Foreman J, Penfield S, Imaizumi T, Millar AJ, et al. 2015. Linked circadian outputs control elongation growth and flowering in response to photoperiod and temperature. Molecular systems biology 11: 776.

Shen Y, Khanna R, Carle CM, Quail PH. 2007. Phytochrome induces rapid PIF5 phosphorylation and degradation in response to red-light activation. Plant physiology 145: 1043–1051.

Shin J, Sánchez-Villarreal A, Davis AM, D. S-X, Berendzen KW, Koncz C, Ding Z, Li C, Davis SJ. 2017. The metabolic sensor AKIN10 modulates the Arabidopsis circadian clock in a light-dependent manner. Plant, cell & environment 40: 997–1008.

Song Y, Jiang Y, Kuai B, Li L. 2018. CIRCADIAN CLOCK-ASSOCIATED 1 Inhibits Leaf Senescence in Arabidopsis. Frontiers in plant science 9: 280.

Souček P, Pavlů J, Medveďová Z, Reinöhl V, Brzobohatý B. 2017. Stability of housekeeping gene expression in Arabidopsis thaliana seedlings under differing macronutrient and hormonal conditions. Journal of plant biochemistry and biotechnology 26: 415–424.

Streb S, Zeeman SC. 2012. Starch metabolism in Arabidopsis. The Arabidopsis book 10: e0160.

Thompson K, Grime JP, Mason G. 1977. Seed germination in response to diurnal fluctuations of temperature. Nature 267: 147–149.

Tsai AY-L, Gazzarrini S. 2012. AKIN10 and FUSCA3 interact to control lateral organ development and phase transitions in Arabidopsis: Interaction between FUSCA3 and AKIN10. The Plant journal: for cell and molecular biology 69: 809–821.

Uhrig RG, Echevarría-Zomeño S, Schlapfer P, Grossmann J, Roschitzki B, Koerber N, Fiorani F, Gruissem W. 2021. Diurnal dynamics of the Arabidopsis rosette proteome and phosphoproteome. Plant, cell & environment 44: 821–841.

Vandesompele J, De Preter K, Pattyn F, Poppe B, Van Roy N, De Paepe A, Speleman F. 2002. Accurate normalization of real-time quantitative RT-PCR data by geometric averaging of multiple internal control genes. Genome biology 3: RESEARCH0034.

Vong GYW, McCarthy K, Claydon W, Davis SJ, Redmond EJ, Ezer D. 2024. AraLeTA: An Arabidopsis leaf expression atlas across diurnal and developmental scales. Plant physiology 195: 1941–1953.

Wang F, Han T, Jeffrey Chen Z. 2024. Circadian and photoperiodic regulation of the vegetative to reproductive transition in plants. Communications biology 7: 579.

Webb AAR, Seki M, Satake A, Caldana C. 2019. Continuous dynamic adjustment of the plant circadian oscillator. Nature communications 10: 550.

Wendell M, Ripel L, Lee Y, Rognli OA, Torre S, Olsen JE. 2017. Thermoperiodic control of floral induction involves modulation of the diurnal FLOWERING LOCUS T expression pattern. Plant & cell physiology 58: 466–477.

Wu G, Anafi RC, Hughes ME, Kornacker K, Hogenesch JB. 2016. MetaCycle: an integrated R package to evaluate periodicity in large scale data. Bioinformatics 32: 3351–3353.

Xiong L, Zhou W, Mas P. 2022. Illuminating the Arabidopsis circadian epigenome: Dynamics of histone acetylation and deacetylation. Current opinion in plant biology 69: 102268.

Xu X, Yuan L, Yang X, Zhang X, Wang L, Xie Q. 2022. Circadian clock in plants: Linking timing to fitness. Journal of integrative plant biology 64: 792–811.

Yeh Y-H, Chang Y-H, Huang P-Y, Huang J-B, Zimmerli L. 2015. Enhanced Arabidopsis pattern-triggered immunity by overexpression of cysteine-rich receptor-like kinases. Frontiers in plant science 6: 322.

Zhang K, Halitschke R, Yin C, Liu C-J, Gan S-S. 2013. Salicylic acid 3-hydroxylase regulates Arabidopsis leaf longevity by mediating salicylic acid catabolism. Proceedings of the National Academy of Sciences of the United States of America 110: 14807–14812.

