## Supplemental Materials for "Stable and dynamic gene expression patterns over diurnal and developmental timescales in *Arabidopsis thaliana*"

Running title: The Circadian Clock in the Ageing Plant


- Ethan J Redmond[1] (https://orcid.org/0000-0001-9105-0609)
- James Ronald[1] (https://orcid.org/0000-0002-8847-0378)
- Seth J Davis[1] (https://orcid.org/0000-0001-5928-9046)
- Daphne Ezer[1,*] (https://orcid.org/0000-0002-1685-6909)

[1]: Department of Biology, University of York, Wentworth Way, Heslington, York YO10 5DD, UK


### Summary of Supplemental Data

#### Table S1 – Gene expression data

This file contains (as separate sheets):

- the normalised gene expression outputs (as transcripts per million; TPM) of all genes, after expression of multiple isoforms of the same gene had been summed
- the median TPM of each set of replicates per condition
- the TPMs of all replicates, for genes which passed filtering for low gene expression

#### Table S2 – GO overrepresentation of top and bottom 10% genes on PC1

This file contains the results from GO term overrepresentation from gProfiler (g:GOSt) (Kolberg *et al.*, 2023) for the genes with the top 10% and bottom 10% loadings on PC1. Columns specific to each set of genes are indicated by the suffixes '_PC1_high' and '_PC1_low' for the top 10% and bottom 10%, respectively.

#### Table S3 – GO overrepresentation of KIN10 target clusters

Like Table S2, this contains GO term overrepresentation results for two clusters of each of the "INCREASED" and "DECREASED" targets from Table S3 from ref. (Baena-González *et al.*, 2007). Clusters match those presented in Figure 2(b) (for "INCREASED" targets) and 2(c) (for "DECREASED" targets).

**Commented [ER1]:** Maybe update the main text to include these terms rather than induced or repressed

#### Table S4 – Results from JTK_cycle showing which genes are rhythmic

Results from JTK_cycle, provided by the wrapper package MetaCycle (v1.2.0) (Hughes *et al.*, 2010; Wu *et al.*, 2016). The JTK_cycle algorithm was run separately on each of the three days – including all biological replicates except those sampled at ZT1, since JTK_cycle requires evenly spaced timepoints. The columns are:

- CycID = the ID of the gene
- BH.Q = the q-value. A gene was classified as rhythmic if this value was less than 0.05.
- ADJ.P = the adjusted p-value
- PER = the predicted period of the gene
- LAG = the predicted phase of the gene
- AMP = the predicted amplitude of the gene

**Commented [ER2]:** Get better info on the stats here

#### Table S5 – Clusters of CCA1 and ELF3 targets

Includes the cluster allocations of predicted targets of CCA1 and ELF3, matching Figure 3(b) and Figure 4(b).

#### Table S6 – GO overrepresentation of CCA1 and ELF3 target clusters

Like Table S2, but the inputs were the cluster allocations from Table S5. There are separate sheets for CCA1 and ELF3 targets and the columns specific to each cluster have the suffixes '_cluster_A', '_cluster_B', etc.

#### Table S7 – Information from CIBERSORTx

The predicted proportions of different cell types from CIBERSORTx (Newman *et al.*, 2019), following the method described in (Vong *et al.*, 2024). The columns are:

- Sample = the ID of the sample
- 17 columns representing different cell types = the proportion of gene expression predicted to come from that cell type for each sample
- P.value
- Correlation
- RMSE

**Commented [ER3]:** Get details from Daphne

#### Table S8 – Results from JTK_cycle showing which cell types are rhythmic

Like Table S4, but with input data from Table S7.

#### Supplemental Figures

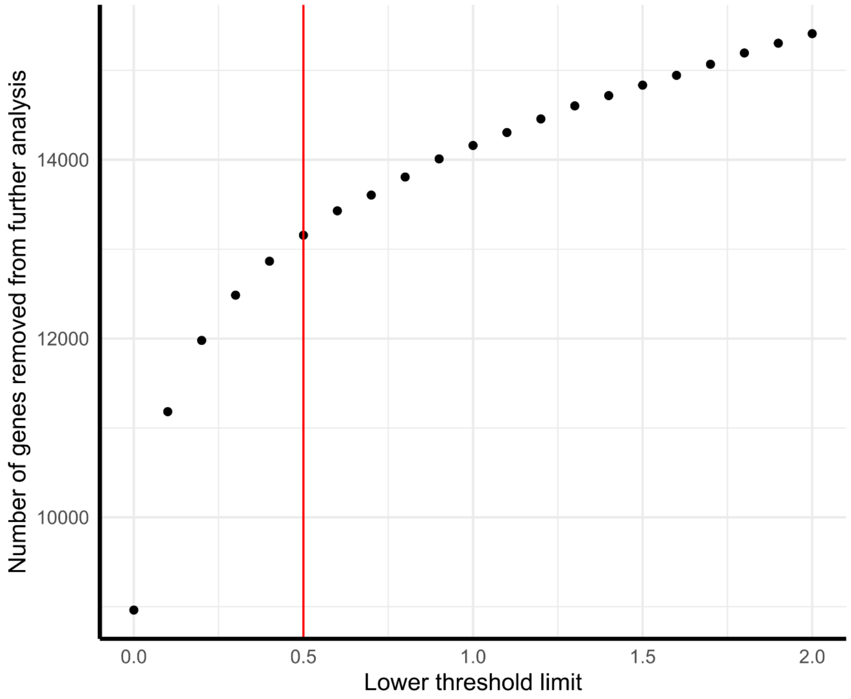


**Supplemental Figure 1.** Genes were filtered out before further analysis if all median TPM values (across the three biological replicates per time-of-day and sampling day) were less than or equal to different 'lower threshold limits'. This figure shows the number of genes removed as the lower threshold limit is increased from 0 to 2, in intervals of 0.1. A final threshold of 0.5 (indicated by the red line) was selected, since the gradient decreases sharply after 0.5.

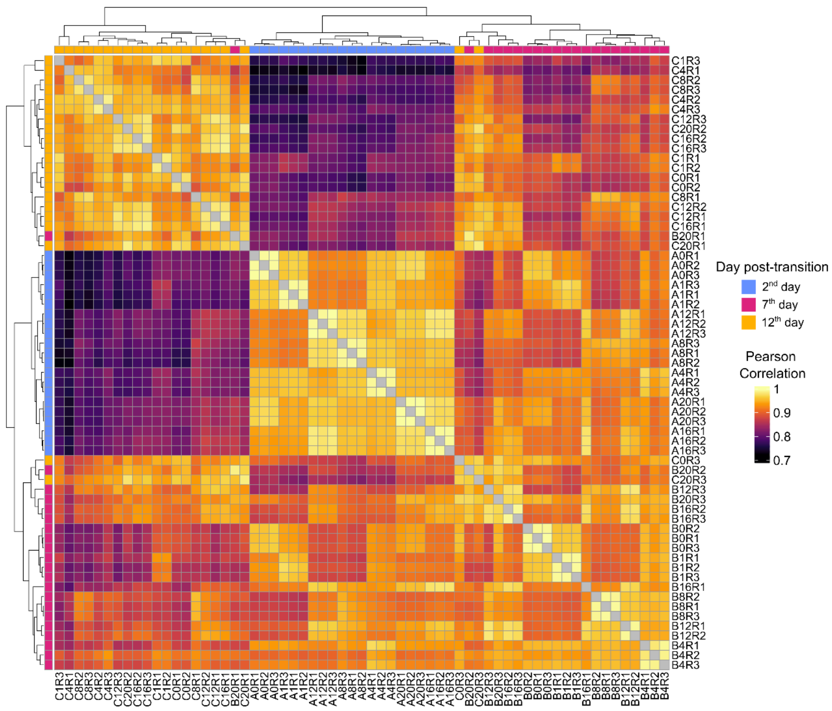


**Supplemental Figure 2.** Pearson **c**orrelation matrix of samples, produced by ComplexHeatmap (version 2.14.0) (Gu *et al.*, 2016). Rows and columns are clustered using hierarchical clustering, using Euclidean distance and the 'complete' method. Most biological replicates cluster together – although 1 sample (B20R1) from the 7th day clustered with samples from the 12th day and 2 samples (C0R3 and C20R3) from the 12th day clustered with samples from the 7th day.

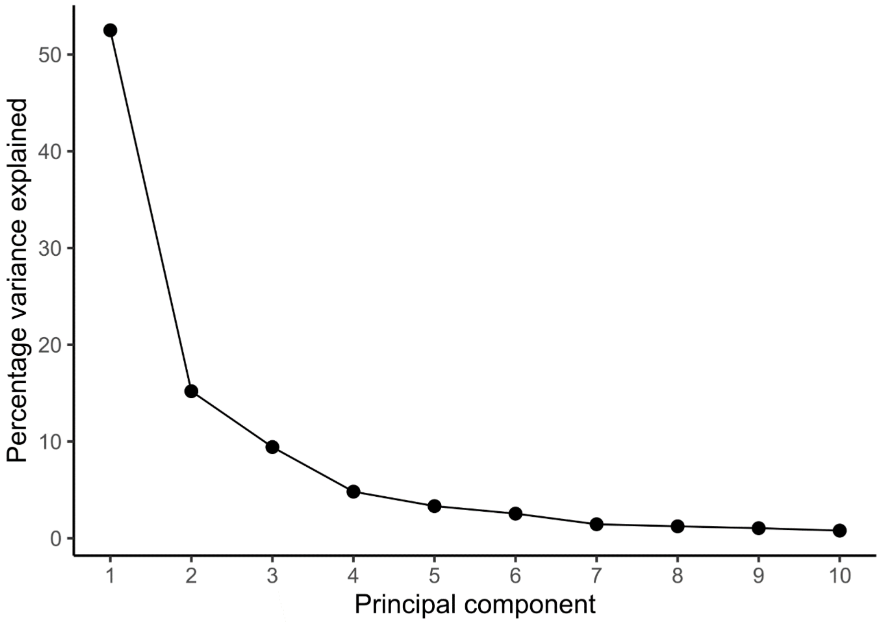


**Supplemental Figure 3.** Scree plot for the first 10 PCs, using log-transformed gene expression as inputs.

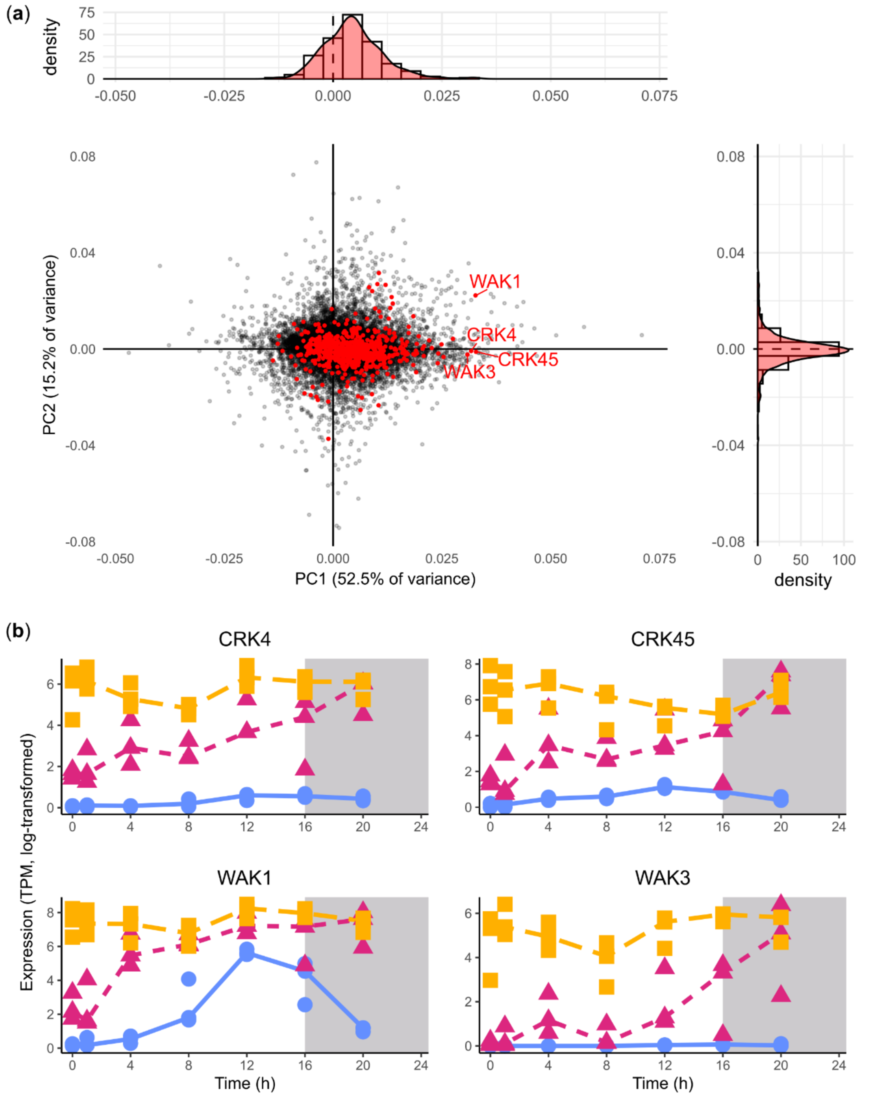


**Supplemental Figure 4. (a)** Loadings of genes on PC1 and PC2, with genes annotated with 'protein serine/threonine kinase activity' (GO:0004674) highlighted in red. The histograms and density plots represent the distributions of the highlighted genes on these axes – scaled so that the area under the density plot is 1. **(b)** Line plots for the 4 genes annotated with 'protein serine/threonine kinase activity' with the highest PC1 loadings. These are also labelled on (a). Points represent individual biological replicates. Lines are drawn through the median value of the biological replicates.

(a)

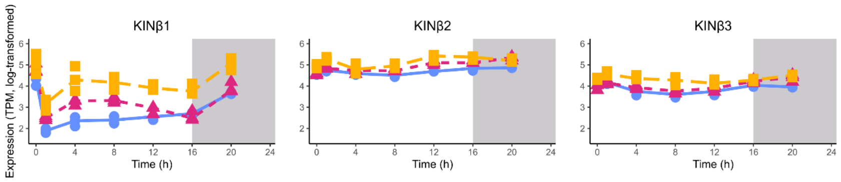


(b)

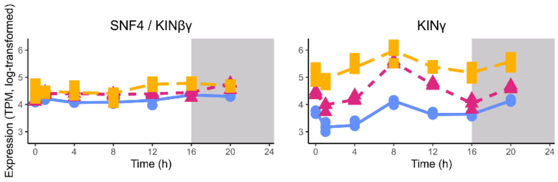


**Supplemental Figure 5. (a)** Expression of the three homologues of the β subunits of SnRK1: KINβ1 (KINBETA1, AT5G21170); KINβ2 (KINBETA2, AT4G16360); and KINβ3 (KINBETA3, AT2G28060). These plots have the same y-axis so expression can be compared between them. **(b)** KINγ (KINgamma, AT3G48530); KINβγ (KINbeta-gamma, HOMOLOG OF YEAST SUCROSE NONFERMENTING 4, SNF4, AT1G09020). These plots have the same y-axis so expression can be compared between them.

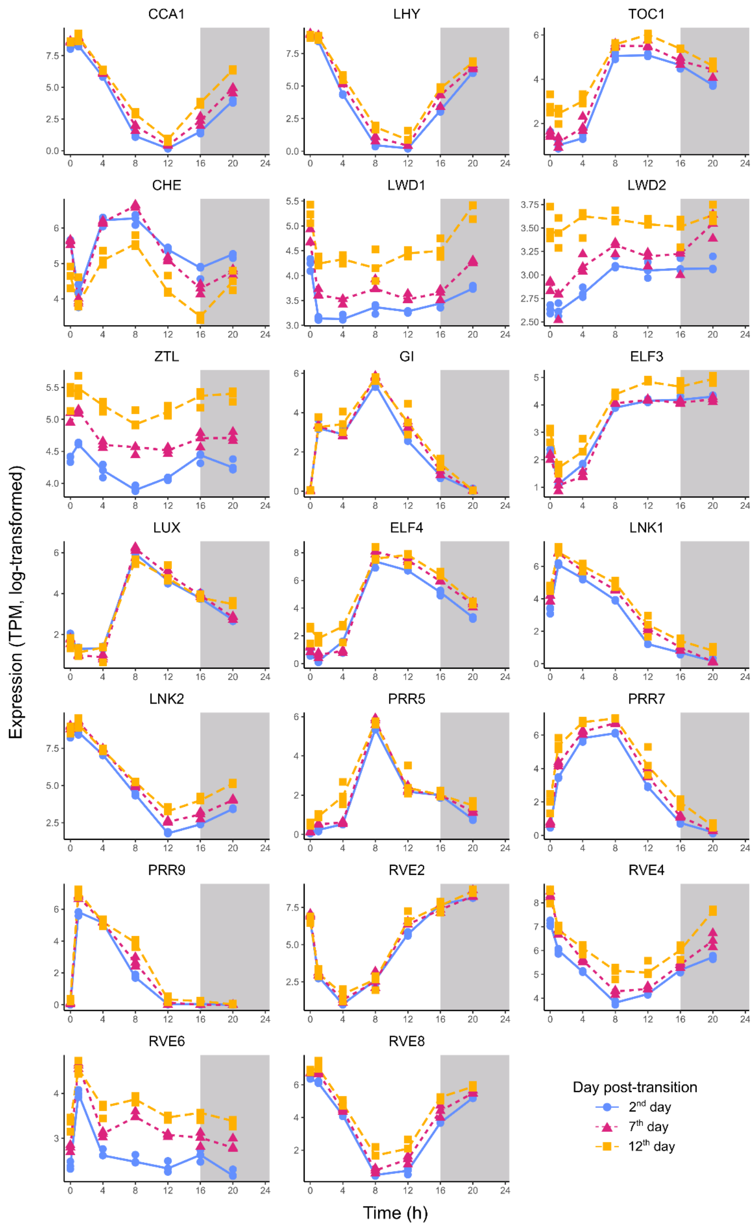


**Supplemental Figure 6.** Expression of all core clock genes, as defined in (Wang *et al.*, 2024).

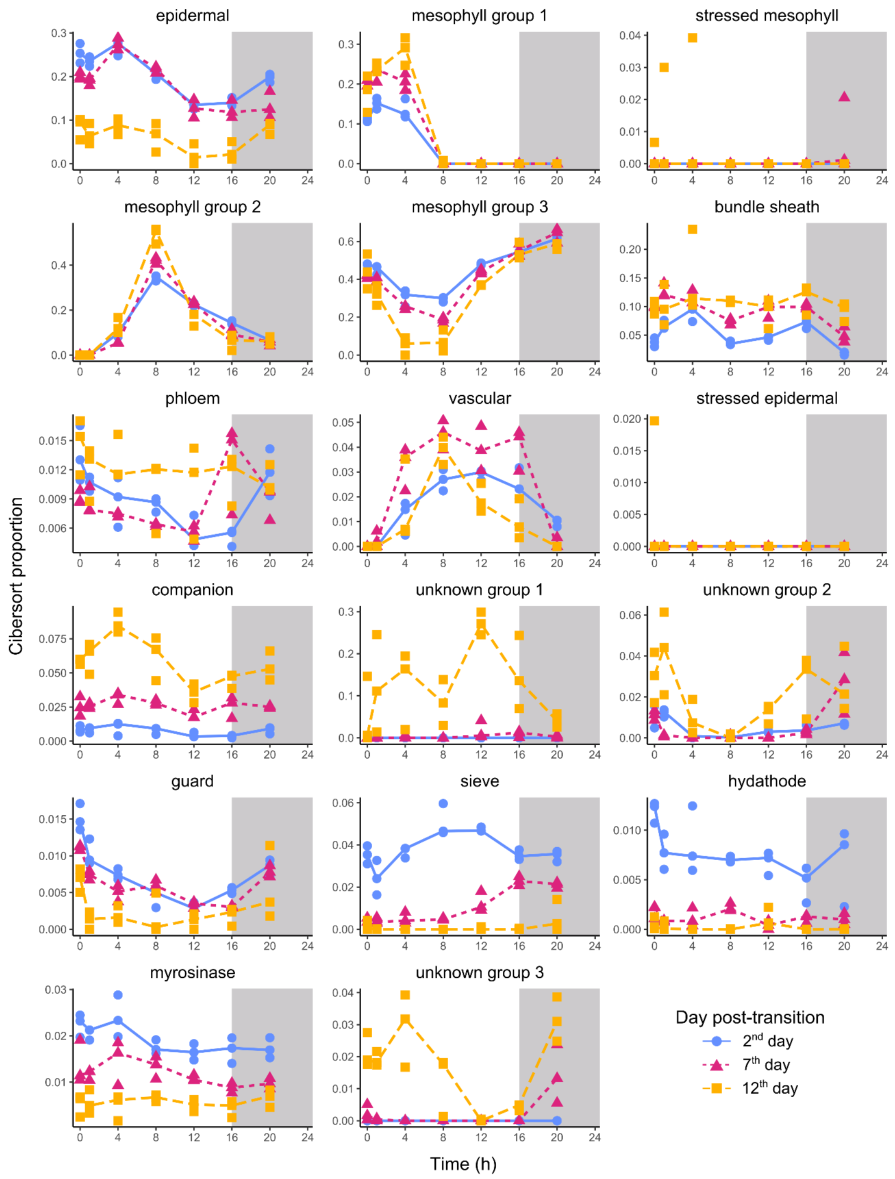


**Supplemental Figure 7.** Predicted proportions of different cell types across all time points, from CIBERSORTx (Newman *et al.*, 2019). Lines are through the medians per timepoint, and points represent individual replicates.

### References


**Baena-González E**, **Rolland F**, **Thevelein JM**, **Sheen J**. **2007**. A central integrator of transcription networks in plant stress and energy signalling. *Nature* **448**: 938–942.

**Gu Z**, **Eils R**, **Schlesner M**. **2016**. Complex heatmaps reveal patterns and correlations in multidimensional genomic data. *Bioinformatics* **32**: 2847–2849.

**Hughes ME**, **Hogenesch JB**, **Kornacker K**. **2010**. JTK_CYCLE: an efficient nonparametric algorithm for detecting rhythmic components in genome-scale data sets. *Journal of biological rhythms* **25**: 372–380.

**Kolberg L**, **Raudvere U**, **Kuzmin I**, **Adler P**, **Vilo J**, **Peterson H**. **2023**. g:Profiler-interoperable web service for functional enrichment analysis and gene identifier mapping (2023 update). *Nucleic acids research* **51**: W207–W212.

**Newman AM**, **Steen CB**, **Liu CL**, **Gentles AJ**, **Chaudhuri AA**, **Scherer F**, **Khodadoust MS**, **Esfahani MS**, **Luca BA**, **Steiner D**, *et al.* **2019**. Determining cell type abundance and expression from bulk tissues with digital cytometry. *Nature biotechnology* **37**: 773–782.

**Vong GYW**, **McCarthy K**, **Claydon W**, **Davis SJ**, **Redmond EJ**, **Ezer D**. **2024**. AraLeTA: An Arabidopsis leaf expression atlas across diurnal and developmental scales. *Plant physiology* **195**: 1941–1953.

**Wang F**, **Han T**, **Jeffrey Chen Z**. **2024**. Circadian and photoperiodic regulation of the vegetative to reproductive transition in plants. *Communications biology* **7**: 579.

**Wu G**, **Anafi RC**, **Hughes ME**, **Kornacker K**, **Hogenesch JB**. **2016**. MetaCycle: an integrated R package to evaluate periodicity in large scale data. *Bioinformatics* **32**: 3351–3353.
